# A nesprin-4/kinesin-1 cargo model for nuclear positioning in cochlear outer hair cells

**DOI:** 10.1101/2021.10.10.463824

**Authors:** Shahar Taiber, Oren Gozlan, Roie Cohen, Leonardo R. Andrade, Ellen F. Gregory, Daniel A. Starr, Yehu Moran, Rebecca Hipp, Matthew W. Kelley, Uri Manor, David Sprinzak, Karen B. Avraham

## Abstract

Nuclear positioning is important for the functionality of many cell types and is mediated by interactions of cytoskeletal elements and nucleoskeleton proteins. Nesprin proteins, part of the linker of nucleoskeleton and cytoskeleton (LINC) complex, have been shown to participate in nuclear positioning in multiple cell types. Outer hair cells (OHCs) in the inner ear are specialized sensory epithelial cells that utilize somatic electromotility to amplify auditory signals in the cochlea. Recently, nesprin-4 (encoded by *Syne4*) was shown to play a crucial role in nuclear positioning in OHCs. *Syne4* deficiency in humans and mice leads to mislocalization of the OHC nuclei and cell death resulting in deafness. However, it is unknown how nesprin-4 mediates the position of the nucleus, and which other molecular components are involved in this process. Here, we show that the interaction of nesprin-4 and the microtubule motor kinesin-1 is mediated by a conserved 4 amino-acid motif. Using in-vivo AAV gene delivery, we show that this interaction is critical for nuclear positioning and hearing in mice. Nuclear mislocalization and cell death of OHCs coincide with the onset of hearing and electromotility and are solely restricted to outer, but not inner, hair cells. Likewise, the *C. elegans* functional homolog of nesprin-4, UNC-83, uses a similar motif to mediate interactions between migrating nuclei and kinesin-1. Overall, our results suggest that OHCs require unique cellular machinery for proper nuclear positioning at the onset of electromotility. This machinery relies on the interaction between nesprin-4 and kinesin-1 motors supporting a microtubule cargo model for nuclear positioning.

## Introduction

Nuclear positioning and movement are integral to cell division, migration and function. Cells have evolved a plethora of mechanisms by which internal and external forces are transmitted to the nucleus, allowing its movement and positioning. The linker of nucleoskeleton and cytoskeleton (LINC) complex is a highly conserved complex of nuclear envelope proteins that has been implicated in nuclear positioning across species and cell types. The LINC complex consists of two families of proteins; SUN (Sad1, UNC-84) proteins that reside in the inner nuclear membrane (INM) and KASH proteins that reside in the outer nuclear membrane (ONM)^1^.

In mammals, five KASH proteins have been identified to date, nesprin-1, nesprin-2, nesprin-3, nesprin-4, and KASH5^2–6^. Mammalian nesprin proteins are comprised of a C-terminal KASH domain and cytoplasmic spectrin repeats (SRs). The KASH domain spans the ONM to the perinuclear space between the INM and ONM, where it interacts with SUN1 and SUN2 proteins that in turn cross the perinuclear space and INM to interact with the nuclear lamina^1,7^. Importantly, localization of nesprins to the nuclear envelope (NE) is dependent on the KASH domain^1,7^.

Unlike nesprin-1 and nesprin-2, with their “giant” isoforms of up to 1014 kDa, nesprin-4 is only 44 kDa, showing conservation of the C-terminal KASH domain but little resemblance to the cytoplasmic regions of nesprin-1 and nesprin-2^5^. Interestingly, expression of *Syne4* is very sparse, and nesprin-4 deficiency in humans and mice was associated with no phenotype other than deafness^8^. In *Syne4*^*-/-*^ mice, nuclei of cochlear outer hair cells are dislocated from their basal position, which is quickly followed by OHC death^8,9^. It was also shown that nesprin-4 interacts with kinesin-1, yet the functional role of this interaction remains unclear^5^. A 4-amino acid motif that mediates binding to the light chain of kinesin-1 in proteins^10,11^ was recently identified in nesprin-2 as well. This binding is critical for nuclear positioning in myotubes ^12^. The authors were able to identify this motif in nesprin-4 as well and speculated that it would mediate the interaction of nesprin-4 with kinesin-1.

In the cochlea, expression of *Syne4* is detected in the two types of mechanosensitive hair cells, inner hair cells (IHC) and outer hair cells (OHC)^8,9^. IHC receive 95% of the afferent cochlear innervation and are responsible for translating sound vibrations to neuronal signals^13,14^. OHC have a unique role in amplifying basement membrane deflection by actively changing their length in response to changes in membrane potential, a behavior termed electromotility. By doing so, OHC contribute to the sensitivity and sharp frequency tuning of the auditory system^15^. OHC are unique to mammals and unique molecular machinery has evolved to enable their specialized function^16^. The discrepancy in the effect of *Syne4* deficiency on IHC and OHC suggests a role for *Syne4* in electromotility.

In this study we aimed at understanding whether OHC nuclear positioning relies on the interaction of nesprin-4 with kinesin-1. We showed the interaction of nesprin-4 and kinesin-1 is mediated by the same 4 amino-acid motif in nesprin-4, as in nesprin-2, and that this interaction is essential for OHC survival. We used AAV gene delivery to test the role of nesprin-4/kinesin-1 interaction *in vivo* and showed that OHC nuclear positioning seems to be entirely dependent on this interaction. We further show that UNC-83, a functional homologue of nesprin-4 in *C. elegans*, interacts with kinesin-1 via the same motif and that disrupting it produces a nuclear migration defect. Finally, examining the evolutionary development of nesprin-4 and the timeline of phenotype onset suggests that the OHC phenotype observed in *Syne4*^*-/-*^ mice could be triggered by electromotility.

## Results

### A conserved motif mediates the interaction of nesprin-4 with kinesin motors

Earlier work identified a conserved kinesin-1 light chain interaction domain, LEWD, in nesprins and showed that it is required for the interaction with kinesin-1^12^. To test the hypothesis that the nucleus in OHC is positioned via kinesin-1 dependent cargo trafficking, we mutated the LEWD kinesin-binding domain in nesprin-4. We introduced a 2-amino acid substitution in this motif at amino acids 243-246, LEWD to LEAA. We then transfected HEK293 cells with a plasmid encoding a FLAG-tagged version of either Nesp4^WT^ (the WT version), Nesp4^AA^ (including the WD>AA mutation), or an empty control. Lysates were subjected to co-immunoprecipitation using anti-FLAG beads and westerns were probed with an anti Kif5b antibody. The results of this experiment showed that this mutation prevents the interaction of nesprin-4 and Kif5b (Figure 1A-C) indicating that the LEWD domain is required for this interaction. In addition, lysates not subjected to immunoprecipitation showed similar band intensities indicating that nesprin-4^AA^ is stable in cultured cells (Supplementary Figure 1A). To test the effect of mutating the LEWD interaction domain on the cellular distribution of nesprin-4, we transfected Chinese hamster ovary (CHO) cells with the same plasmids encoding both versions of nesprin-4 fused to FLAG or an empty FLAG control. We found that nesprin-4^AA^ localized to the nuclear envelope similarly to nesprin-4^WT^. Hence, nesprin-4^AA^ was properly translated and trafficked to the nuclear envelope despite being unable to bind kinesin-1 (Figure 1D-E).

**Figure 1.**
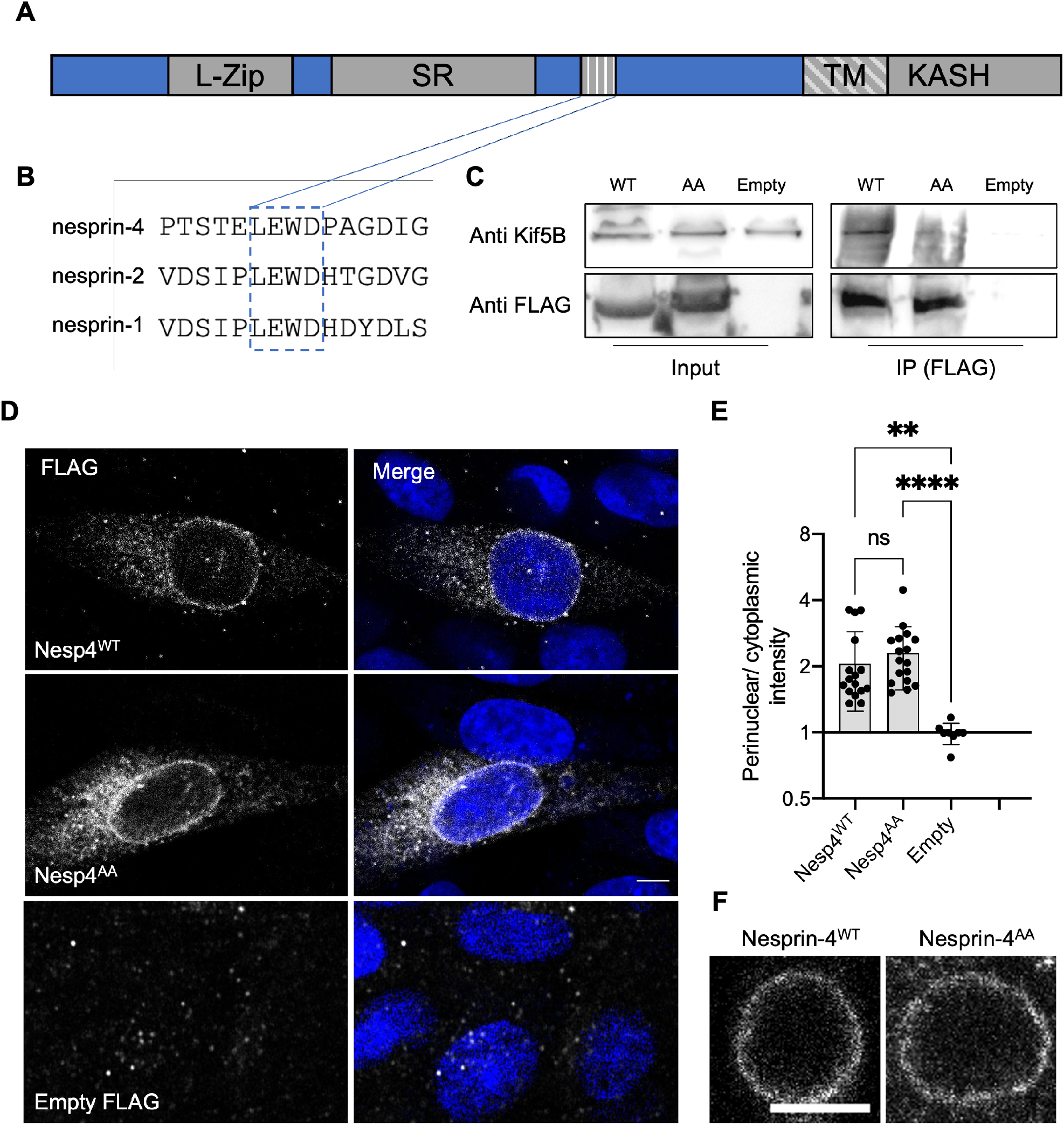
LEWD motif in mouse nesprin-4 is required for kinesin-1 mediated nuclear migration. A. Schematic of nesprin-4 sequence showing the LEWD motif and other domains: TM - transmembrane, SR - spectrin repeat, L-Zip - leucine zipper. B. Alignment of human nesprin-4, nesprin-2, and nesprin-1 LEWD motifs. C. Co-immunoprecipitation of LEWD and LEAA nesprin-4 with kif5b. Input = total lysate without precipitation. D. Nesprin-4^WT^ and nesprin-4^AA^ are recruited to the nuclear envelope. Nesprin-4 was labeled by FLAG (gray) and nuclei by DAPI (blue). E. Quantification of nuclear recruitment shows that WT nesprin-4 and mutant nesprin-4^AA^ are localized to the NE similarly, n=16 cells for Nesp4^WT^, n=17 cells for Nesp4^AA^ and n=8 for empty FLAG, from three independent experiments. F. Zoomed-in Whole-mount immunofluorescence of OHC nuclei from P9 old mice injected with AAV.Syne4^WT^ or AAV.Syne4^AA^ at P1 shows similar localization to the nuclear envelope. Nesprin-4 labeled by FLAG (gray) and nuclei by DAPI (blue). Plots show mean ± SD Statistical test was Kruskal-Wallis test with Dunn’s correction for multiple comparisons. ns=not significant, **P<0.01, ****P<0.0001. Scale bars = 5 μm

To study the effect of mutated nesprin-4 in hair cells, we generated AAV vectors encoding either the wild type or the LEAA variant of nesprin-4 (termed AAV.Syne4^WT^ and AAV.Syne4^AA^, respectively). We used AAV9-PHP.B as we have shown previously that it can transduce IHC and OHC very efficiently when injected during the first 48 h postnatally. Here, we injected mice with AAV.Syne4^AA^ and observed that AAV.Syne4^AA^ transduces OHC efficiently and that nesprin-4^AA^ localizes to the NE of OHC, as we previously showed for WT nesprin-4^9^, further indicating that it was properly translated and localized *in vivo* (Figure 1F, Supplementary Figure 1B).

### Rescue of *Syne4* deafness is kinesin-1 dependent

We have shown before that *Syne4*^*-/-*^ mice show early-onset progressive deafness and that at P14 the nuclei of OHC are dislocated towards the apical surfaces, leading to rapid loss of OHC^8,9^. In addition, we have shown that a single injection of AAV9-PHP.B encoding *Syne4* was sufficient to entirely prevent nuclear mislocalization, OHC loss, and deafness in *Syne4*^*-/-*^ mice. To test whether nesprin-4 function is dependent on the EWD motif, we injected *Syne4*^-/-^ mice with either AAV.Syne4^WT^ or AAV.Syne4^AA^ at P0-P1.5.

We observed that while AAV.Syne4^WT^ rescues nuclear localization in OHC, AAV.Syne4^AA^ fails to do so (Figure 2A-B). Moreover, unlike AAV.Syne4^WT^, AAV.Syne4^AA^ does not rescue OHC survival at 4w and only the OHC at the very apex survive, as in untreated *Syne4*^-/-^ mice (Figure 2C-D, Supplementary Figure 2). In contrast to OHC, the position of IHC nuclei was not affected in the *Syne4*^*-/-*^ mice. Auditory brainstem response (ABR) of *Syne4*^*-/-*^ mice injected with AAV.Syne4^WT^ was markedly rescued at 4w with some treated *Syne4*^-/-^ showing thresholds indistinguishable from WT (Figure 2E). However, *Syne4*^-/-^ mice injected with AAV.Syne4^AA^ were not significantly different from untreated *Syne4*^-/-^ mice. Distortion-product otoacoustic emissions (DPOAE) are generated by the OHC electromotility-dependent cochlear amplifier, and thus we measured DPOAEs as an assay to assess OHC functionality *in vivo*. In line with the ABR recordings, AAV.Syne4^AA^ did not rescue DPOAE thresholds, while AAV.Syne4^WT^ fully rescued thresholds in *Syne4*^-/-^ mice (Figure 2F). Thus, the LEWD interaction domain in nesprin-4 is required for proper nuclear positioning and survival of OHC.

**Figure 2.**
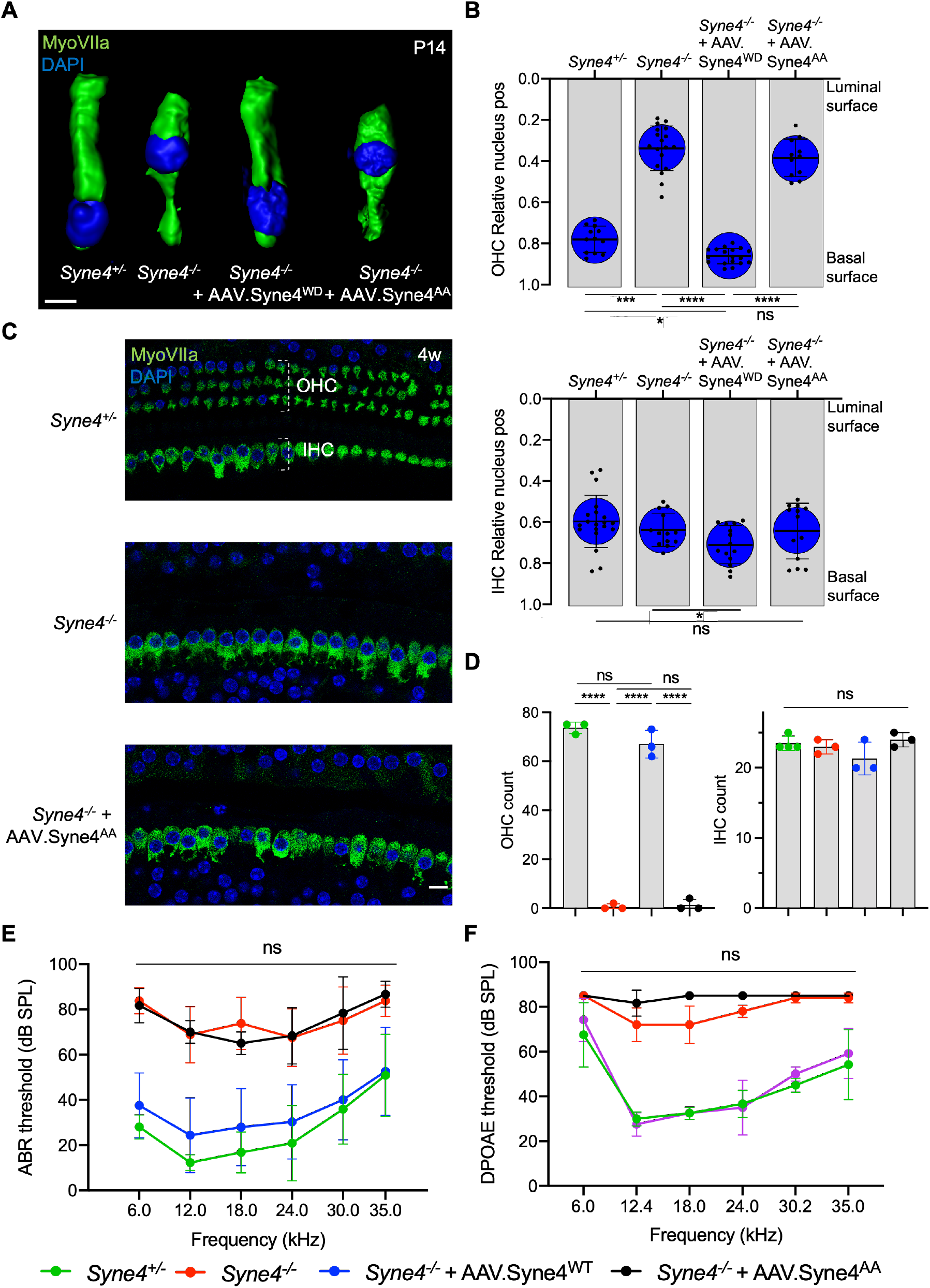
Mutation of the nesprin-4 LEWD motif abolishes the activity of nesprin-4 in the inner ear. A. 3D surface projection of P14 OHC from the 12kHz region of *Syne4*^*+/+*^, *Syne4*^*-/-*^, *Syne4*^*-/-*^ injected with AAV.Syne4^WT^, and *Syne4*^*-/-*^ injected with AAV.Syne4^AA^ mice. B. Image analysis quantification of relative nucleus position at P14 for OHC and IHC. A total of 12 OHC and 20 IHC from 2 *Syne4*^*+/-*^ mice, 18 OHC and 13 IHC from 3 *Syne4*^*-/-*^ mice, 18 OHC and 14 IHC from 3 *Syne4*^*-/-*^ mice injected with AAV.Syne4^WT^, and 11 OHC and 13 IHC from 2 *Syne4*^*-/-*^ mice injected with AAV.Syne4^AA^ were analyzed. C. Whole-mount immunofluorescence of the 12kHz region from 4w old *Syne4*^*+/+*^, *Syne4*^*-/-*^, and *Syne4*^*-/-*^ mice injected with AAV.Syne4^AA^. Hair cells labeled by MyoVIIa (green) and nuclei by DAPI (blue). D. OHC and IHC counts at the 12kHz region per 200 µm. E. ABR thresholds of 4w old *Syne4*^*+/+*^, *Syne4*^*-/-*^, *Syne4*^*-/-*^ injected with AAV.Syne4WT, and *Syne4*^*-/-*^ injected with AAV.Syne4AA mice, n=11 for *Syne4*^*+/+*^, n=8 for *Syne4*^*-/-*^, n=20 for *Syne4*^*-/-*^ + AAV.Syne4^WT^, n=3 for *Syne4*^*-/-*^ + AAV.Syne4^AA^. F. DPOAE thresholds at 4w, n=6 for *Syne4*^*+/+*^, n=5 for *Syne4*^*-/-*^, n=6 for *Syne4*^*-/-*^ + AAV.Syne4^WT^, n=3 for *Syne4*^*-/-*^ + AAV.Syne4^AA^. Plots show mean ± SD. Statistical tests were Kruskal-Wallis with Dunn’s correction for multiple comparisons for B, one-way ANOVA for D and 2-way ANOVA for E and F with Tukey’s correction for multiple comparisons. ns=not significant, *P<0.05, ***P<0.001, ****P<0.0001. Scale bars = 10 µm.

### Nesprin-4^AA^ acts as a dominant regulator of nesprin-4 activity

To verify that the inability of nesprin-4^AA^ to rescue OHC loss and hearing is not due to reduced transduction efficiency or compromised protein stability, we tested whether overexpression of nesprin-4^AA^ will recapitulate the knockout phenotype by competing with the endogenous nesprin-4 for binding to limited SUN1 sites in the nuclear envelope. To test this we injected *Syne4*^+/-^ mice (that have no phenotype) with either version of nesprin-4. While overexpression of nesprin-4^WT^ had no observable effect on OHC nuclear position at P14 or OHC survival at 4w, overexpression of nesprin-4^AA^ leads to mislocalization of OHC nuclei and OHC loss (Figure 3A-D). Consistent with our previous observation, IHC nuclear position and IHC survival were not affected (Figure 3B-D). ABR results of *Syne4*^+/-^ mice injected with AAV.Syne4^AA^ showed only moderate levels of hearing loss, probably because only one ear was injected (Figure 3E). ABR electrodes were positioned to measure the injected ear, but it is known that there is contribution of both ears to the recorded ABR signal^17^. In addition to the contribution of the contralateral ear, it is possible that surviving OHC in the apical region of injected cochleae preserve auditory sensitivity of low frequencies. DPOAE results measured from the injected ear showed a much clearer result, as massive loss of OHC in *Syne4*^+/-^ mice injected with AAV.Syne4^AA^ abolished cochlear amplification (Figure 3F).

**Figure 3.**
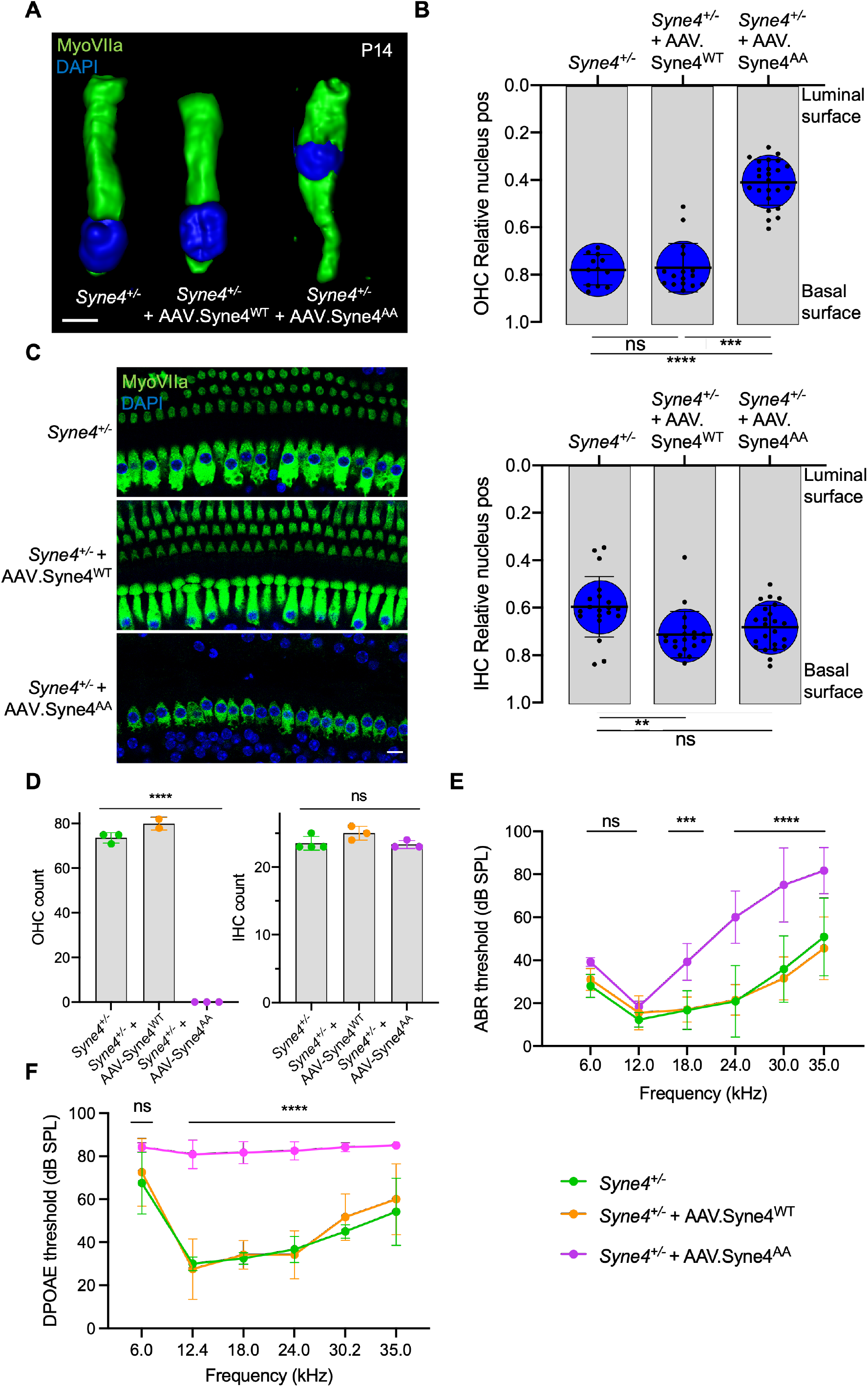
Overexpression of mutant nesprin-4^AA^ has a dominant negative effect on OHC. A. 3D surface projection of P14 OHC from the 12kHz region of *Syne4*^*+/-*^, *Syne4*^*+/-*^ injected with AAV.Syne4^WT^, and *Syne4*^*+/-*^ injected with AAV.Syne4^AA^ mice. B. Image analysis quantification of relative nucleus position at P14 for OHC and IHC. A total of 12 OHC and 20 IHC from 2 *Syne4*^*+/-*^ mice, 16 OHC and 19 IHC from 4 *Syne4*^*+/-*^ mice injected with AAV.Syne4^WT^, and 24 OHC and 23 IHC from 4 *Syne4*^*+/-*^ mice injected with AAV.Syne4^AA^ were measured. C. Whole-mount immunofluorescence of the 12kHz region from 4w old *Syne4*^*+/-*^ and *Syne4*^*+/-*^ injected with AAV.Syne4^AA^ mice. Hair cells labeled by MyoVIIa (green) and nuclei by DAPI (blue). D. OHC and IHC counts at the 12kHz region per 200 um. E. ABR thresholds of 4w old *Syne4*^*+/+*^, *Syne4*^*+/-*^ injected with AAV.Syne4^WT^, and *Syne4*^*+/-*^ injected with AAV.Syne4^AA^ mice, n=11 for *Syne4*^*+/+*^, n=10 for *Syne4*^*+/-*^ + AAV.Syne4^WT,^ and n=6 for *Syne4*^*+/-*^ + AAV.Syne4^AA^. F. DPOAE thresholds at 4w, n=6 for *Syne4*^*+/+*^, n=6 for *Syne4*^*+/-*^ + AAV.Syne4^WT^, and n=6 for *Syne4*^*+/-*^ + AAV.Syne4^AA^. Plots show mean ± SD. Statistical tests were Kruskal-Wallis with Dunn’s correction for multiple comparisons for B, one-way ANOVA for D and 2-way ANOVA for E and F with Tukey’s correction for multiple comparisons. ns=not significant, **P<0.01, ***P>0.001, ****P<0.0001. Scale bars = 10 µm.

### Disrupting the EWD motif of a nesprin-4 homologue in *C. elegans* produces a nuclear migration defect

To further define the role of the LEWD motif in nesprin-4 function *in vivo*, we made use of a functional homologue of nesprin-4 in *C. elegans*, UNC-83. UNC-83 is an outer nuclear membrane KASH protein that interacts with the kinesin light chain, KLC-2, to recruit kinesin-1 heavy chain, UNC-116, to the surface of nuclei. Kinesin-1 then provides the force to move nuclei toward the plus ends of microtubules in embryonic hypodermal (hyp7) precursor cells^18,19^. Mutations in *unc-83, klc-2*, or *unc-116* lead to a failure of nuclear migration in hyp7 cells^18,19^. We hypothesized that UNC-83 interacts with kinesin-1 via the same conserved EWD motif we identified in nesprin-4 (located here at residues 331-333). We mutated the EWD motif to GSA and assayed nuclear migration. The *unc-83(GSA)* mutant protein localized normally to the nuclear envelope (Figure 4A), suggesting that disrupting the EWD motif of the endogenous protein does not compromise protein stability or localization *in vivo*. However, there was a significant hyp7 nuclear migration defect in *unc-83(GSA)* similar to the *unc-83(null)* defect^20^, assayed by counting mislocalized hyp7 nuclei in the dorsal cord (Figure 4B). In larval P cells, UNC-83 mediates nuclear migration through the minus-end-directed motor dynein, and following nuclear migration, P cells differentiate into vulval cells and GABA neurons^21,22^. Missing GABA neurons suggest that P-cell nuclear migration was unsuccessful. The *unc-83(GSA)* mutant had no significant effect on P-cell nuclear migration, suggesting that the mutant protein retains its dynein-related functions (Figure 4C). Together, these data suggest that disruption of the EWD motif in both nesprin-4 and UNC-83 leads to the production of a stable protein that is properly folded and localized, but unable to function through kinesin-1 to mediate nuclear positioning. Moreover, the conserved role of the EWD motif in mice and C. elegans suggests that the interaction between kinesin and nesprin is associated with conserved functions in vivo.

**Figure 4.**
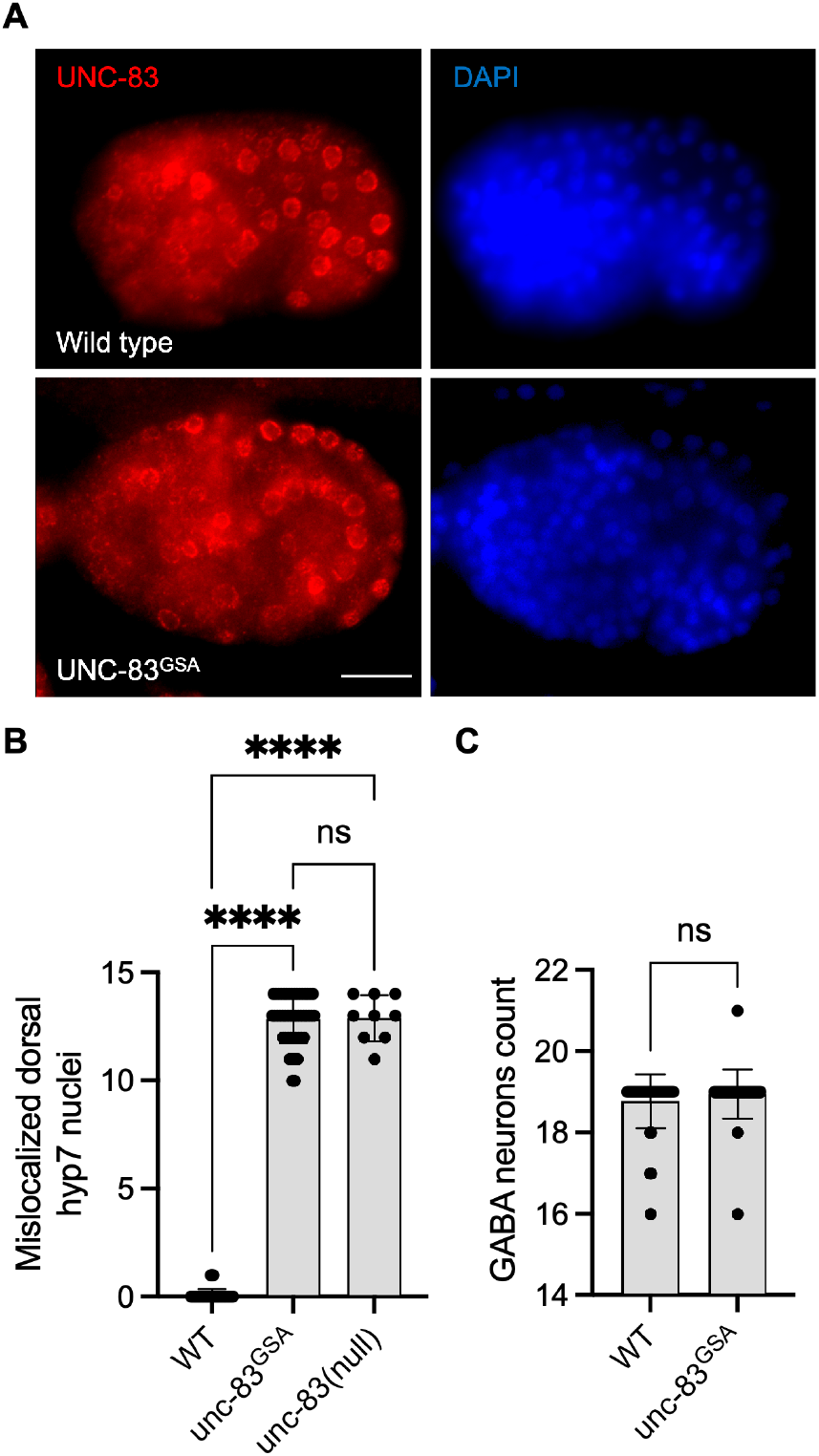
Mutation of the EWD motif of UNC-83, the functional homolog of nesprin-4, disrupts kinesin-1 mediated nuclear migration. A. Mutant *unc-83(GSA)* (red) localized to the nuclear envelope, similar to WT UNC-83 (DAPI-stained nuclei are blue) in *C. elegans* bean-stage embryos. Dorsal is up and anterior is left. B. Hypodermal nuclear migration defects were observed in *unc-83(null)* and *unc-83(GSA)* but not in wild-type embryos, n=50 for wild type, n=9 for *unc-83(null)*, and n=48 for *unc-83(GSA)*. C. GABA neuron counts in *unc-83(+)* and *unc-83(GSA)* embryos are similar, n=40 for each group. Plots show mean ± SD. Statistical test was Kruskal-Wallis test with Dunn’s correction for multiple comparisons for B and Mann-Whitney for C. ns=not significant, ****P<0.0001. Scale bar = 5 µm

### *Syne4*^*-/-*^ OHC phenotype coincides with the onset of electromotility and hearing

To understand why *Syne4* deficiency leads to OHC death we investigated the temporal dynamics of *Syne4* expression with respect to the onset of the phenotype. Single molecule fluorescent in situ hybridization (smFISH) performed at E16, P1 and P11 shows that the *Syne4* is expressed by both IHC and OHC at early stages of development, in agreement with previous analyses we performed on transcriptome data (Figure 5A)^9^. We then analyzed the position of the nucleus in *Syne4*^*-/-*^ mice at ages P8, P10, P12, and P14 using confocal immunofluorescence and observed that while *Syne4* is expressed early in embryonic development, nuclei are mislocated in *Syne4*^*-/-*^ OHC only at P12-P14, coinciding with the onset of OHC electromotility and hearing (Figure 5B)^23^: at P8 there is no difference in nuclear position between *Syne4*^*-/-*^ and *Syne4*^*+/-*^ OHC and by P12 the nuclei of *Syne4*^*-/-*^ OHC is significantly mislocalized.

**Figure 5.**
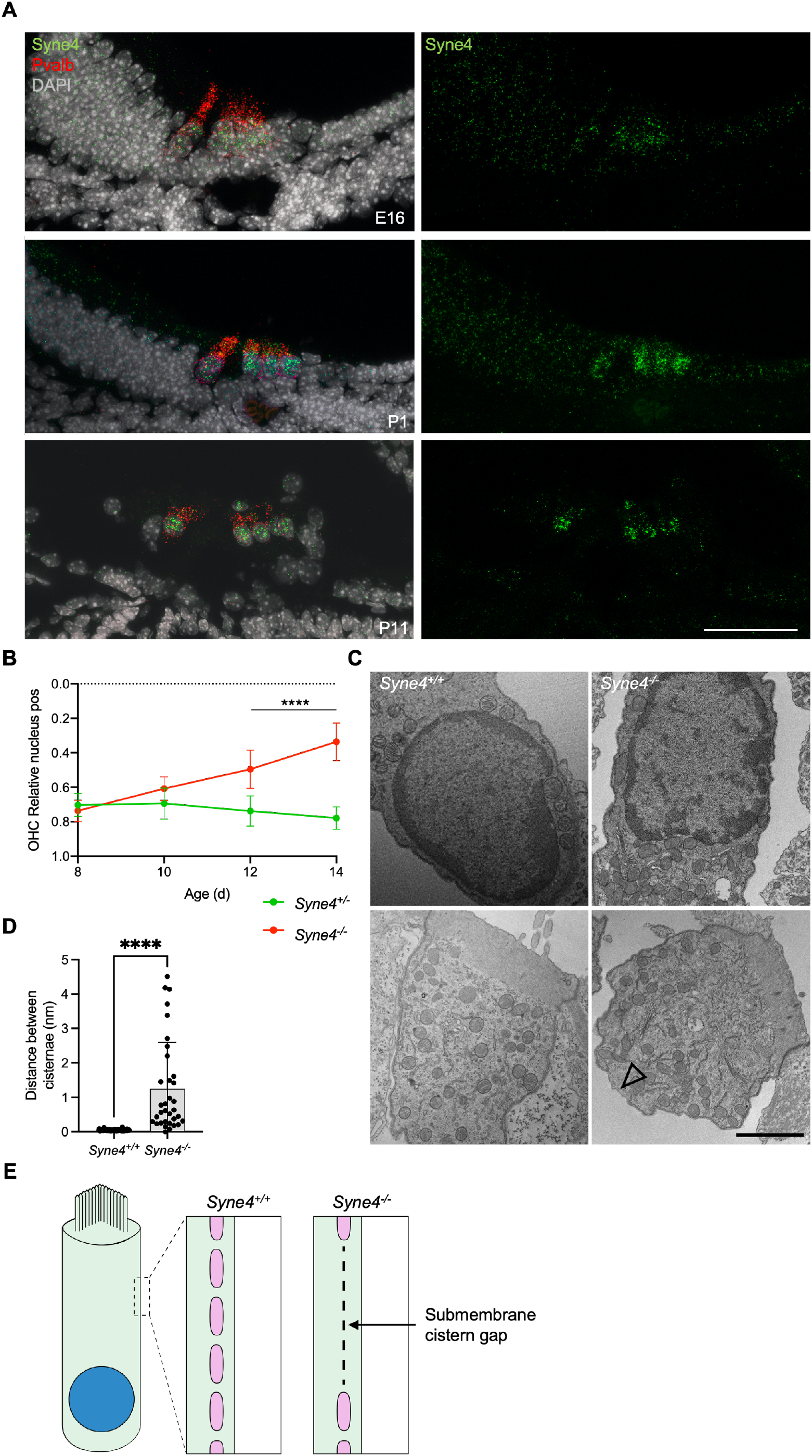
*Syne4*^*-/-*^ phenotype is correlated to onset of electromotility and hearing. A. smFISH for *Syne4* (green) and Pvalb (red) performed at E16, P1, and P11. Nuclei stained with DAPI (gray). B. Image analysis quantification of relative nucleus position at P8-P14 for OHC in *Syne4*^*-/-*^ and *Syne4*^*+/-*^ mice. A total of 39 OHC from 8 *Syne4*^*+/-*^ mice, and 64 OHC from 11 *Syne4*^*-/-*^ mice were measured. C. TEM of P11 *Syne4*^*-/-*^ and *Syne4*^*+/+*^ mice. Open arrow denotes sub-membrane cisternae defect. Experiment was repeated 3 times. D. Quantification of distance between submembrane cisternae. A total of 7 cells from 5 Syne*4*^*+/+*^ mice, and 10 cells from 2 *Syne4*^*-/-*^ mice were measured. E. Schematic illustration of the observed cisternae defects. Plot shows mean ± SD. Statistical test was multiple t-test with two-stage step-up correction for multiple comparisons for B and Mann-Whitney for D. ****P<0.0001. Scale bars = 50 µm for A, 2 µm for C.

### Cellular defects in *Syne4*^*-/-*^ OHC

To study the ultrastructural effects of nucleus mislocalization in *Syne4*^*-/-*^ OHC we performed TEM on P12 *Syne4*^*+/+*^ and *Syne4*^*-/-*^ ears. We observed no obvious differences in the inner architecture of the nucleus (Figure 4C). In both *Syne4*^*-/-*^ and *Syne4*^*+/+*^ OHC nuclei the nuclear envelope was continuous, with no blebs or ruptures, and heterochromatin was similarly localized to the nuclear periphery. Overall, we did not observe any clear defects in sub-nuclear architecture. Interestingly, we did observe defects in the sub-membrane cisternae of *Syne4*^*-/-*^ OHCs that were not detected in *Syne4*^*+/+*^ OHCs (Figure 5C and Supplementary Figure 3). The sub-membrane cisternae in OHCs are membranous structures consisting of ordered particles found beneath the lateral plasma membrane of OHCs^24^. Their role is currently unknown, but they have been suggested to contribute to the mechanical or electrical properties of OHCs^25^. To quantify this observation, we measured the lateral distance between adjacent submembrane cisternae (i.e., the length of the gaps between them) in *Syne4*^*-/-*^ and *Syne4*^*+/+*^ OHC and found a significant increase in the mean distance cisternae in *Syne4*^*-/-*^ OHC (Figure 5D-E). Together these results may suggest that loss of nesprin-4 causes damage to the intricate structure of the lateral wall of OHCs. More experiments are required to determine the mechanism leading to these defects.

### Evolution of nesprin-4

OHC are unique to mammals and have specialized proteins and structures that support their electromotility function. Considering the time of onset of nuclear dislocation, and the specificity to OHC, we analyzed the evolution of the nesprin protein family to understand how nesprin-4 emerged. Sequences of *Homo sapiens* and *Mus musculus* nesprin proteins were aligned against homologous sequences from chicken *(Gallus gallus)*, zebrafish *(Danio rerio)*, and frog *(Xenopus tropicalis)*. The results were used to build a phylogenetic tree of nesprin proteins (Supplementary Figure 4). Nesprin-4 from *Homo sapiens* and *Mus musculus*, and *Xenopus tropicalis* cluster with a *Danio rerio* protein named “uncharacterized protein LOC777613”. Alignment of its sequence with the *Homo sapiens* and *Mus musculus* nesprin-4 paralogs suggest that this protein is in fact the zebrafish homolog of nesprin-4 (Figure 6A). Inspection of the expression of this protein in published datasets via gEAR shows enrichment in HC^26^ (Figure 6B-C). No hit was detected for nesprin-4 or the KASH domain of nesprin-4 in the chicken genome. We note that the extensive sequence divergence of the cytoplasmic portions of KASH proteins limits the strength of BLAST analyses to relatively close species.

**Figure 6.**
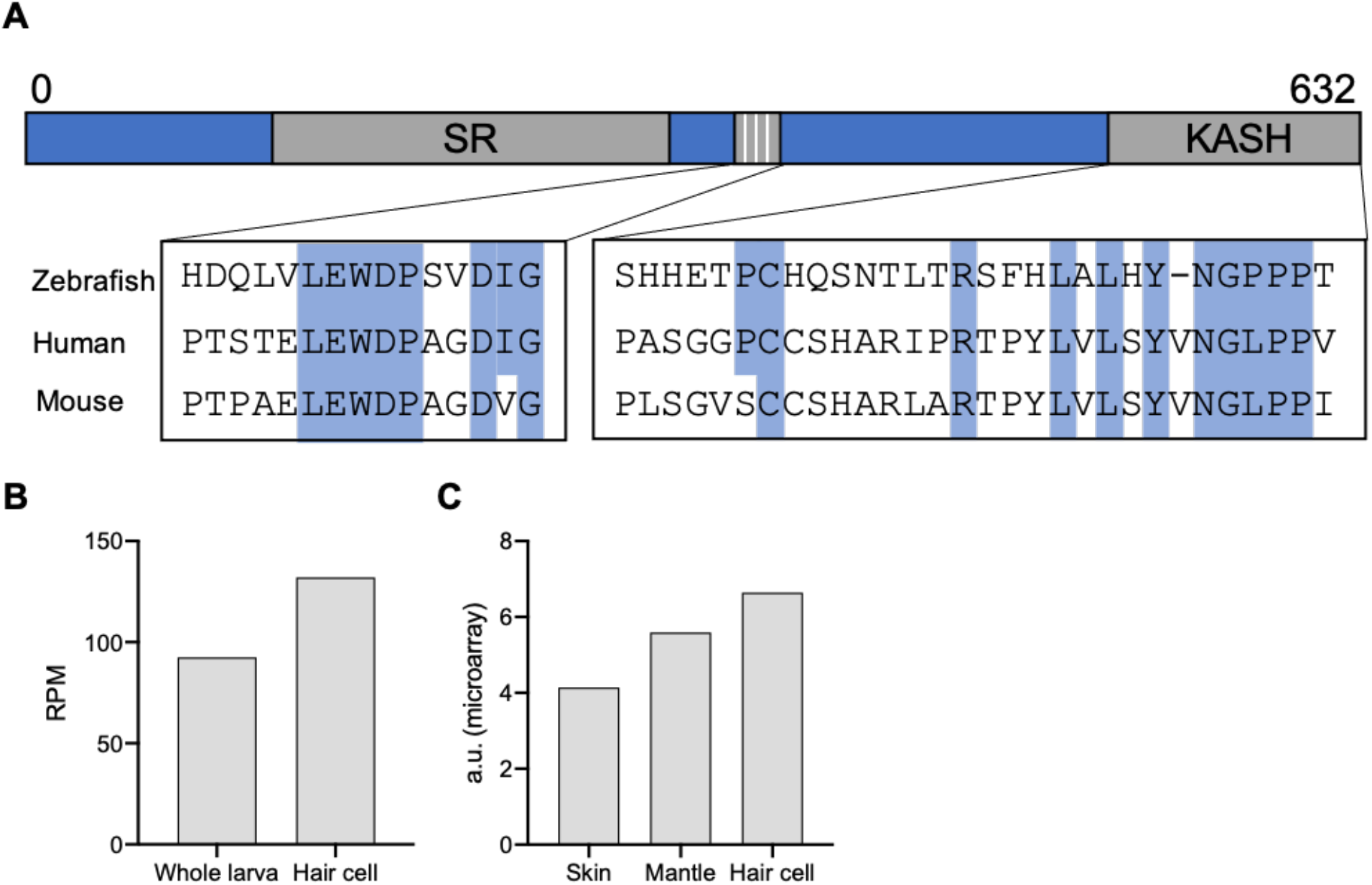
Identification of a new putative zebrafish nesprin-4. A. Alignment of the putative zebrafish nesprin-4 protein with human and mouse nesprin-4 proteins. The nuclear KASH domain and cytoplasmic kinesin-1 binding motif are highlighted. SR = spectrin repeat. B. Expression of putative zebrafish nesprin-4 by RiboTag RNAseq^27^.. C. Expression of putative zebrafish nesprin-4 by microarray^28^. Data curated using gEAR^26^.

## Discussion

Here, we provide evidence supporting a cargo model of nuclear positioning in OHC, dependent on the interaction between nesprin-4 and kinesin-1. We used co-immunoprecipitation to show that nesprin-4 interacts with kinesin-1 through a conserved 4 amino acid motif, LEWD. Using *in vivo* AAV gene delivery, we demonstrate that disruption of this motif creates an inactive nesprin-4 and recapitulates the *Syne4*^*-/-*^ hearing loss phenotype in mice. Using gene editing, we show that this motif in a functional homologue of nesprin-4 in *C. elegans* is required for the migration of nuclei in hypodermal cells. We show that the cellular phenotype in *Syne4*^*-/-*^ mice is not correlated to the expression of *Syne4*, but rather to the onset of electromotility and hearing. Finally, we suggest a nesprin-4 ortholog in zebrafish.

Kinesin-1 is composed of a dimer of kinesin heavy chains and two kinesin light chains (KLCs). A recent study showed that Klc2 deficient mice exhibit hearing loss but no nuclear positioning defect in HC, which would argue against the role of kinesin-1 in nuclear positioning in cochlear HC^29^. However, as other KLCs exist in mammals, we suggest that Klc2 is redundant for nuclear positioning but required for another cellular function, such as phospholipid processing, as suggested by the authors.

We show that while delivery of WT *Syne4* by AAV is sufficient to prevent nuclear mislocalization, OHC death, and hearing loss, delivery of *Syne4* in which the kinesin-binding motif has been mutated (*Syne4*^*AA*^) leads to no rescue whatsoever. Moreover, AAV.Syne4^AA^ recreates the *Syne4*^*-/-*^ phenotype in *Syne4*^*+/-*^ mice. This effect can be interpreted as dominant-negative, in which nesprin-4^AA^ competes with endogenous nesprin-4 to bind SUN1, thus preventing a functional nucleus-cytoskeleton interaction. In line with these findings, we show that mutating the EWD motif in UNC-83, a *C. elegans* homologue of nesprin-4, causes a nuclear migration defect. These results suggest that the interaction of nesprin-4 with kinesin-1 is conserved to mediate nuclear localization both in the mammalian inner ear as wells as *C. elegans* hypodermal cells.

The fact that we did not observe (both in this work and in^8,9^) any effect on IHC nuclear position or survival is intriguing. Why is nesprin-4 so critical for OHC function and survival but not for IHC? Our data show that the onset of the phenotype in OHC correlates with the onset of electromotility. We show that while *Syne4* is expressed as early as E16, there is no difference in nuclear position until around P12-P14. Non-linear membrane capacitance, used as a proxy of electromotility, cannot be detected before P9 and continues to increase until P17-18^23^. DPOAE measurements, which are used as an *in vivo* functional assessment of OHC function, are first identified at P11 for 8kHz and P13 for 30kHz, with growing amplitudes over the next two weeks^30^. Together, these observations suggest that at the time of phenotype onset in *Syne4*^*-/-*^ mice, the organ of Corti should be subjected to vertical pulling and pushing forces associated with OHC electromotility (Figure 7). These observations therefore support the hypothesis that nesprin-4/kinesin-1 nuclear positioning mechanism is necessary for holding the nucleus in place at the onset of somatic electromotility that is restricted to OHCs.

**Figure 7.**
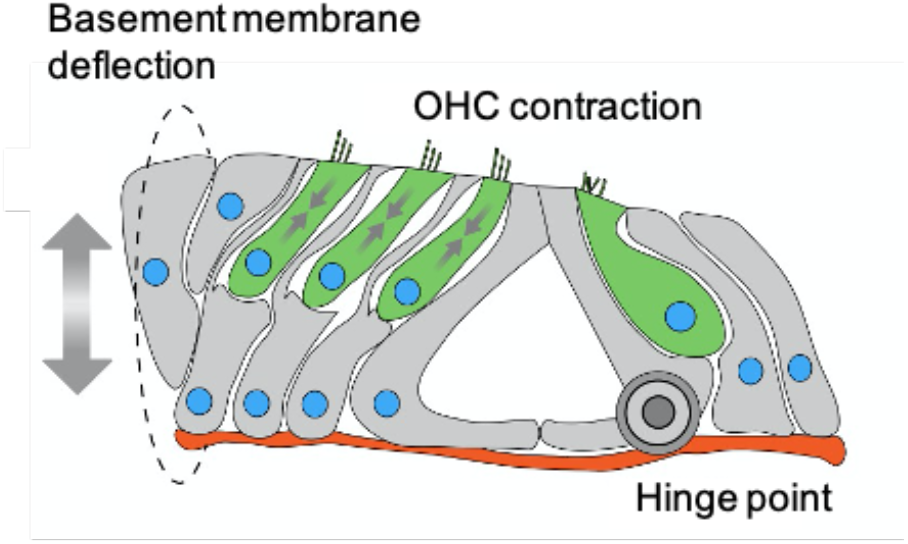
Schematic model depicting mechanical forces applied on OHC nuclei of mature organ of Corti.

It is interesting to note that IHCs express other nesprins as well, such as nesprin-1 and nesprin-2^26^. These nesprins are expressed at lower levels in IHC compared to nesprin-4 but, as for the case of nesprin-4, their expression patterns are similar to that of OHCs^26^. This observation suggests that perhaps low levels of other nesprins are sufficient to position the nucleus in OHCs in early stages but are not sufficient to hold it once electromotility starts.

Our results are consistent with a kinesin-1 dependent cargo model of nuclear positioning. While other mechanisms of nuclear positioning have been demonstrated in other cell types, such as anti-parallel sliding microtubules in myotubes^31^ and dynein-mediated transport In the developing eye of drosophila^32^, these are likely not sufficient for adult OHC.

It remains unclear why nuclear mislocalization leads to OHC death. The changes in subcellular organization we observe could impair cellular function and damage organelles. In addition, mechanical strain on nuclei has been shown to lead to NE rupture and DNA damage, with LINC complex proteins implicated in several cell types^33^. Furthermore, the LINC complex can regulate gene expression via interaction with chromatin. Intriguingly, our TEM data did not reveal any clear change in the structure of the NE or in the organization of the chromatin, yet submembrane cisternae defects were detected in the lateral wall of *Syne4*^*-/-*^ OHC. This suggests that loss of nesprin-4 causes cellular defects outside the nucleus. Further investigation of these aspects could elucidate the important roles of nesprin-4 and the interplay between mechanical forces and organelle structure and positioning in maintaining cellular homeostasis.

While IHCs are very well conserved, appearing in all vertebrates, OHCs, with their distinct behavior of electromotility, are unique to mammals^35^. Many key OHC genes can be found in close relatives of mammals in the vertebrate lineage, with evidence showing the adaptation they underwent to enable the emergence of this highly specialized cell. A classic example of this principle is prestin, the motor of electromotility and a canonical OHC marker, that is ubiquitous in chordates^36^. Vestibular hair cells express prestin but do not exhibit electromotility^37^. Spectrin beta-V, which is part of the lateral wall cytoskeletal complex, has also undergone evolutionary changes to support the emergence of electromotility^38^.

Our evolutionary analysis suggests that the zebrafish uncharacterized protein LOC777613 is a homolog of mammalian nesprin-4. Examination of the expression of this gene in published datasets reveals that it is probably upregulated in hair cells. *Xenopus tropicalis* has a likely nesprin-4 homolog as well (named nesprin-4), but no expression or localization data are available at this point. As zebrafish hair cells are not electromotile, investigating the role of the putative zebrafish nesprin-4 could put to test our hypothesis regarding the role of electromotility in this phenotype.

## Materials and methods

### Mice

All animal procedures were approved by the Animal Care and Use Committee (IACUC) at Tel Aviv University (01-17-101) and performed according to the NIH Guide for the Care and Use of Laboratory Animals. For every experiment, the age is specified in the figure legend. Mice were maintained on a C57Bl/6J background. Genotyping was performed on DNA prepared from ear punch biopsies, and extracted and amplified using the KAPA HotStartMouse Genotyping Kit (Sigma, KK7352). Genotyping primers for the WT allele were WT_FWD (5-ACTCCCAGCTCCAAGCTACA-3) and WT_REV (5-GCAGAGCCAAAGAAACCAAG-3), and for the galactosidase gene were LacZ_FWD (5-GTCTCGTTGCTGCATAAACC-3) and LacZ_REV (5-TCGTCTGCTCATCCATGACC-3). Cycling conditions were an initial 3-min denaturation at 95°C followed by 35 cycles of 30 s 95°C, 30 s 60°C, and 30 s at 72°C, with a final elongation of 3 min at 72°C. PCR products were loaded into a 2% agarose ethidium-bromide gel for electrophoresis.

### C. elegans

The *unc-83(EWD-GSA)* mutant strain (UD628: *unc-83(yc74)*) was created with CRISPR/Cas9 gene editing of the endogenous *unc-83* locus using a *dpy-10* co-CRISPR approach^39–43^. The *dpy-10* single-stranded DNA repair template and crRNA^39^ were added to a mix containing *unc-83* crRNA (5’–TGCTGCACAAGCCTATGAAT– 3’; Dharmacon), *unc-83 EWD-GSA* single-stranded repair template (5’– CCGGGATAGTGACACGGCGCCAGAACATAGTGATGCTGCACAAGCCTATGGATCCGCTGAGTATAATGTAAGTGTTCAAACTGTTTAATGCAGAAAAAATTGAACTT– 3’; Dharmacon), universal tracrRNA (Dharmacon), and purified Cas9 protein (UC Berkeley QB3). The preassembled Cas9 complex was injected into the gonad of hermaphrodite young adult animals. Offspring were screened for the roller phenotype caused by the *dpy-10(gof)* edit, indicating successful CRISPR/Cas9 action, and their progeny were screened for the *unc-83(EWD-GSA)* mutation by PCR (Forward primer 5’-CGACACATCCTCCCCATTGAAG-3,’ reverse primer 5’-GTGGTTTTAGGAGACTTAGTAGTAC-3’) followed by digestion with BamHI, which only cuts mutant *unc-83(EWD-GSA)* DNA. Homozygous mutant animals were verified by Sanger sequencing. *unc-83(EWD-GSA)* mutants were crossed to integrated, nuclear GFP expressed from a hyp7 promoter^44^ or a GFP GABA neuron marker^45^ to make strains UD643: *unc-83(yc74); ycIs9* and UD660: *unc-83(yc74); oxIs12 [unc-47::GFP]*, respectively. Embryos were fixed and stained with monoclonal antibody 1209D7 against UNC-83c as previously described^20^. Nuclear migration assays were done by counting embryonic hypodermal precursor nuclei that are mislocalized in the dorsal cord, or by counting missing P-cell lineages in the ventral cord as previously described^46^. *C. elegans* were visualized using a Leica DM6000 fluorescence microscope with a 63 × Plan Apo 1.40 NA objective, a Leica DC350 FX camera, and Leica LAS AF software.

### Plasmids and AAV production

pEMTB-3xGFP was a gift from Eran Perlson. The AAV2 plasmid containing the *Syne4* coding sequence was previously generated^9^. To introduce the desired WD to AA substitution, a gene block was synthetized, and restriction ligation cloning was performed. AAV.Syne4^WD^ viral preps were generated as described before^9^. AAV.Syne4^AA^ was produced at Tel Aviv University using AAVpro Purification Kit Maxi (Takara) according to the manufacturer’s instructions. Briefly, 12 15 cm plates of HEK293T cells were triple-transfected with the AAV9-PHP.B cap plasmid, pAd5 adenovirus helper plasmid, and insert (ITR) Syne4^AA^ plasmid. 72 h after transfection cells were harvested and viral vectors extracted. Viral titers were calculated based on qPCR amplification with the following primers: Syne4_FWD (5-cctcttcccatgagcatcaa-3), Syne4_REV (5-ccggaagttcaacctcaaca-3). The AAV2/9.PHP.B.CMV.3xFLAG.Syne4^WD^.bGH titer was 7.7E + 12 gc/ml, and the AAV2/9.PHP.B.CMV.3xFLAG.Syne4^AA^.bGH titer was 1.5737E + 13 gc/ml. Vectors were aliquoted into 10 μl vials and stored at −80°C until use.

### Animal surgery

Animal surgery was performed as described previously^47^. Briefly, A posterior-semicircular canal (PSCC) injection was carried out in mice at P0–P1.5. Mice were anesthetized by induced hypothermia and kept on a cold surface throughout the procedure. Vector solution (1.0–1.2 ul) was aspirated into a borosilicate glass pipette held by a stereotaxic device and connected to a CMA 102 Microdialysis Pump (CMA, Sweden). Once identified, the PSCC was gently punctured and the virus was microinjected for ∼ 2 min (∼ 10 nl/s). After surgery, mice were placed on a heating pad for recovery before being returned to their mothers.

### Auditory testing

ABR and DPOAE, measurements were performed as described previously^47^. Briefly, mice anesthetized by intra-peritoneal injection of a combination of ketamine (100 mg/kg) and xylazine (10 mg/kg). Mice were presented with click stimuli and pure tones at 6, 12, 18, 24, 30, and 35 khz, at intensities ranging from 10 to 90 dB-SPL, in steps of 5 dB. All measurements were performed using an RZ6 multiprocessor, MF1 speakers (Tucker-Davis Technologies, Alachua, FL), and an ER-10b+ microphone (Etymotic Research, Elk Grove Village, IL), and analyzed using BioSigRZ software (Tucker-Davis Technologies, Alachua, FL) and a designated R algorithm (Rstudio, Boston, MA). All experiments were performed by the same tester.

### Cell culture

HEK293 and CHO cells were cultured in Dulbecco’s Modified Eagle’s Medium supplemented with 10% FBS, 1% penicillin, and 1% L-glutamine (Biological Industries) in a humidified incubator at 37°C with 5% CO_2_. HEK293 cells were transfected using homemade PEI and CHO cells were transfected using LT-1 according to the manufacturer’s instructions. For protein localization experiments, CHO cells were co-transfected with either nesprin-4^WT^ or nesprin-4^AA^ plasmids and pEMTB-3xGFP to visualize the cytoplasm.

### Immunofluorescence

Inner ear dissection and staining were performed as previously described^47^. Samples were imaged using a Zeiss LSM. 880 (Zeiss, Oberkochen, Germany). CHO cells were fixed in 5% PFA for 30 min, permeabilized in 0.2% triton for 10 min, and blocked in 5% BSA for 1 h. Antibodies were diluted in 5% BSA for cells and in primary antibody diluent (BarNaor) for inner ear samples. Antibody staining concentrations were as follows: rabbit polyclonal myosin VIIa (Proteus Biosciences, 25-6790) 1:250, rabbit anti-FLAG (Abcam, ab205606) 1:100, DAPI (Abcam ab228549) 1:1,000, and goat anti-rabbit Alexa Fluor 488 (Cell Signaling 4412s) 1:250.

### Image analysis

All data processing was performed off-line using commercial software packages (MATLAB R2019b, MathWorks Inc, Natick, MA, and Fiji). For 3D surface projections, Imaris 8.4 software was used (Bitplane, Belfast, UK). Nuclei position was extracted using a semi-automatic custom-made code in Matlab. For each cell, the positions of the apical surface, nucleus centroid and basal end of the cell were manually marked as reference points. Then, a smooth curve delineating the main axis of the cell was fitted using the marked reference points. The total length of the cell was estimated by calculating the total length of the fitted curve. The nuclear position with respect to the apical surface was estimated as the length of the curve between the apical and nuclear reference points. Nuclear recruitment was quantified in custom-made Fiji macro. Nuclei were segmented using the DAPI channel and auto-thresholding. The entire cell was segmented using the microtubules channel. Cytoplasmic intensity was measured by measuring the intensity in the segment of the entire cell excluding the segment nucleus. Perinuclear intensity was measured in 1.1 µm band surrounding the nucleus.

### Transmission electron-microscopy

Following decapitation and extraction of whole inner ears, 1 mL of fixation solution (2.5 % glutaraldehyde, 4% PFA, 0.1 M sodium cacodylate, 5 mM CaCl2, and 2 mM MgCl2) was slowly injected through the round window and samples were transferred to a 10 mL glass vial with fixative for 2h at RT. After dissection samples were post-fixed in 1% osmium tetroxide with 1.2% potassium ferricyanide for 40 min RT. Samples were then washed 3 times in buffer, stained with 1% uranyl acetate for 1h, dehydrated in graded acetone dilutions till absolute, embedded with Epon resin and polymerized for 2 days at 60°C. 60-nm thickness sections were cut (Leica UC7 ultramicrotome), transferred to copper grids and stained with UranyLess (EMS) for 5 minutes. Samples were imaged using Zeiss Libra TEM at 80 kV.

### Western blot and immunoprecipitation

HEK293 cells were cultured in Dulbecco’s Modified Eagle’s Medium supplemented with 10% FBS, 1% penicillin, and 1% L-glutamine (Biological Industries), transfected with AAV.Syne4^WD^, AAV.Syne4^AA^, or empty FLAG control plasmids, using jetPEI (Polyplus) according to manufacturer’s instructions and harvested 48 h after transfection. One 10 cm plate per condition was lysed 1 ml RIPA buffer (Sigma) and Halt Protease Inhibitor Cocktail (Thermo). Tubes were placed on an end-over-end shaker for 1 h at 4°C followed by centrifugation at 16,000 g for 15 min at 4°C. 20 ul of the resulting supernatant was used as input and the rest was incubated with EZview anti-FLAG M2 beads (F2426, Sigma) for 1-2 h at 4°C on an end-over-end shaker. Beads were then washed and eluted in sample buffer according to manufacturer’s instructions and loaded into a 10% SDS-PAGE gel. Samples were then transferred onto a nitrocellulose membrane which was then blocked in 3% skimmed milk (BD Difco). Membranes were then subjected to immunoblotting. Kif5b was detected by anti Kif5b (Abcam, ab167429). FLAG was detected using rabbit anti DDDDK antibody (Abcam, ab205606). Blots were visualized using anti rabbit HRP antibody (Cell Signaling Technologies, 7074) and SuperSignal West Pico PLUS Chemiluminescence Substrate (Thermo Scientific).

### Evolutionary analysis

The nesprin protein sequences were aligned using MUSCLE^48^. The maximum-likelihood phylogeny was generated using IQ-Tree^49^ with the LG+F+R5 model which was the best fitting model both according to the Bayesian information criterion (BIC) and corrected Akaike information criterion. Support values of the phylogenetic tree are derived from 1,000 ultrafast bootstrap replicates^50^.

### In situ hybridization

Single molecule fluorescent in situ hybridization (smFISH) detection of expression of *Syne4* was performed as described previously^51^. Briefly, cochleae were dissected and fixed in 4% PFA overnight at 4°C. Postnatal day 11 cochleae were decalcified in 0.25 M EDTA overnight. Then, Samples were washed, subjected to a sucrose gradient and embedded in OCT. Next, samples were sectioned on a cryostat at 10 µm thickness. Probes for *Syne4, Pvalb*, a marker of hair cells, and the RNAscope Fluorescent Multiplex Reagent Kit were purchased from Advanced Cell Diagnostics. Hybridization was performed following the manufacturer’s suggested protocol. Results were imaged using a Zeiss LSM 710 confocal microscope.

### Statistics

Statistical tests, group sizes, and P values are noted in the figure legends. Littermates were randomized to the different experiment groups. No blinding was performed, and all tests were carried out by the same tester. Objective measures were preferred when possible. These include hair cell counts, DPOAE thresholds, image analysis, and evolutionary analyses. Statistical analyses were performed using Prism 8 software (GraphPad, San Diego, CA). When required, Shapiro–Wilk and Kolmogorov–Smirnov were used to test data for normality. For comparisons of more than two groups or conditions, the Tukey post hoc test was used to adjust P values when data passed normality tests and Kruskal-Wallis test with Dunn’s correction was used when data did not pass normality tests.

## Data availability

This study includes no data deposited in external repositories. All data and/or codes are available upon request.

## Acknowledgements

The authors wish to thank Daniel Nataf, Adi Barzel, and Michal Lisnyansky Bar-El for technical help and advice. The research was funded by the National Institutes of Health/NIDCD R01DC011835 (K.B.A.) and DC000059 (M.W.K.), the National Institutes of Health/NIGMS R35GM134859 (D.A.S.), the United States-Israel Binational Science Foundation (BSF) 01027150, Jerusalem, Israel (K.B.A.), the Israel Science Foundation (grant no. 1763/20) (K.B.A., M.W.K.), and the European Research Council (ERC) under the European Union’s Horizon 2020 Research and Innovation Program, Grant Agreement No. 682161 (D.S.). K.B.A. is an incumbent of the Drs. Sarah and Felix Dumont Chair for Research of Hearing Disorders. U.M. is supported by the Chan-Zuckerberg Initiative Imaging Scientist Award, NSF NeuroNex Award No. 2014862, and the Hillblom Foundation. U.M. and L.R.A. are supported by National Institutes of Health (NIH) grant no. R21 DC018237, the Waitt Foundation, the Grohne Foundation, and NIH-NCI CCSG: P30 014195. This work was performed in partial fulfillment of the requirements for a Ph.D. degree by Shahar Taiber, recipient of the Klass Family Fellowship, at the Faculty of Medicine, Tel Aviv University, Israel.

## Author contributions

ST, KBA, DS, and DAS designed the study and interpreted the results. ST performed molecular biology experiments, mouse injections, auditory testing, cochlear dissections, and immunofluorescence, and analyzed the data. LA and UM performed electron microscopy and analyzed the results. RH and MWK designed and performed RNAscope experiments. OG and RC performed cell culture experiments and microscopy. YM performed evolutionary analyses. RC wrote MATLAB codes for image analysis. EFG performed all the *C. elegans* experiments. ST, KBA, DS, and DAS wrote the manuscript. All authors contributed to the article and approved the submitted version.

---

**Supplementary Figure 1.**
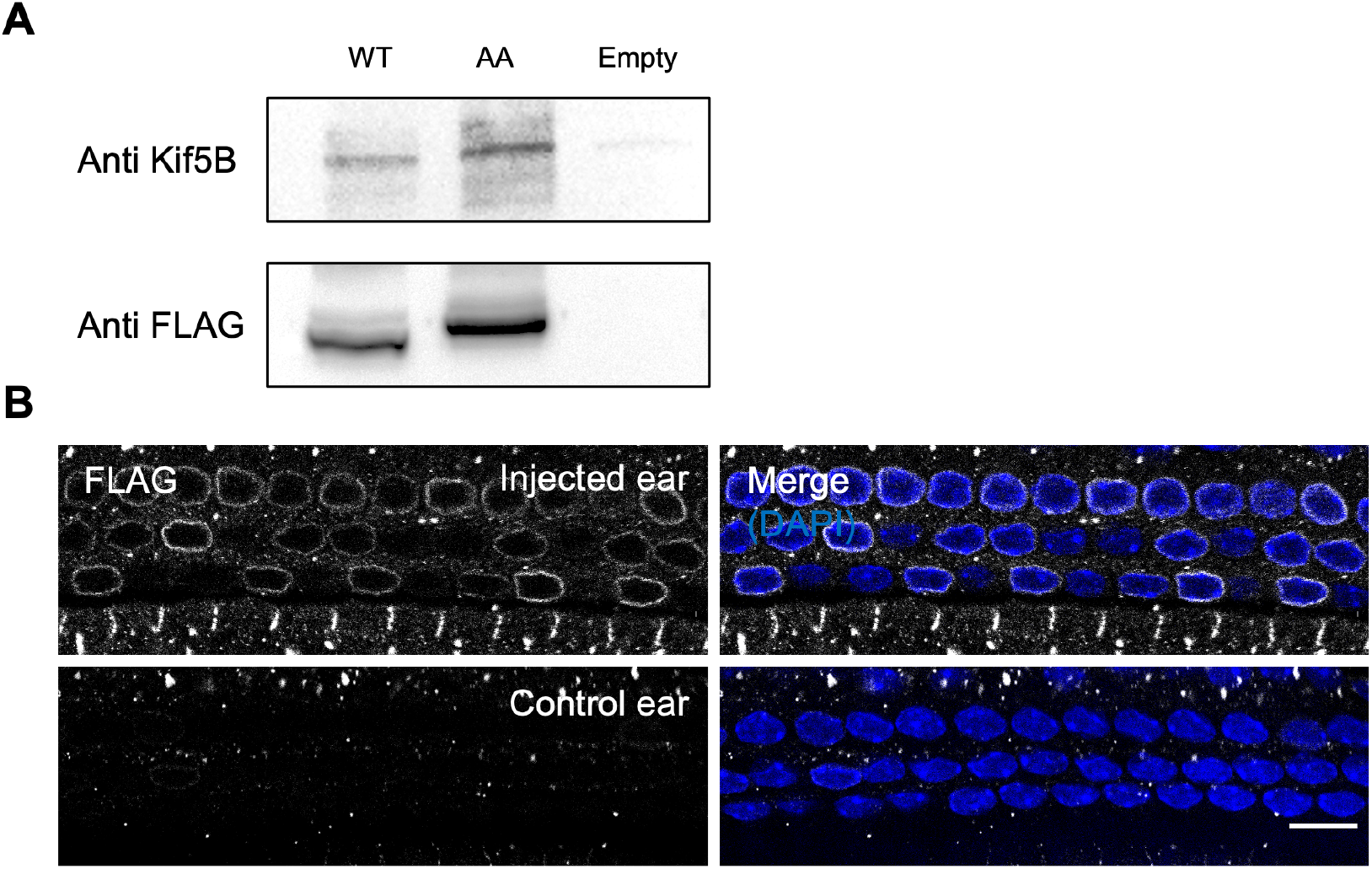
A. Representative western blot performed on protein extracted from transiently transfected HEK293 cells. Nesprin-4^WT^ and nesprin-4^AA^ were detected using an anti-FLAG antibody and kif5b was used as loading control. Similar band intensities indicate that nesprin-4^AA^ is stable in-vivo. B. Whole-mount immunofluorescence from a P9 old mouse injected with AAV.Syne4^AA^ at P1 shows efficient transduction of OHC. Nesprin-4 labeled by FLAG (gray) and nuclei by DAPI (blue). Scale bars = 10 μm.

**Supplementary Figure 2.**
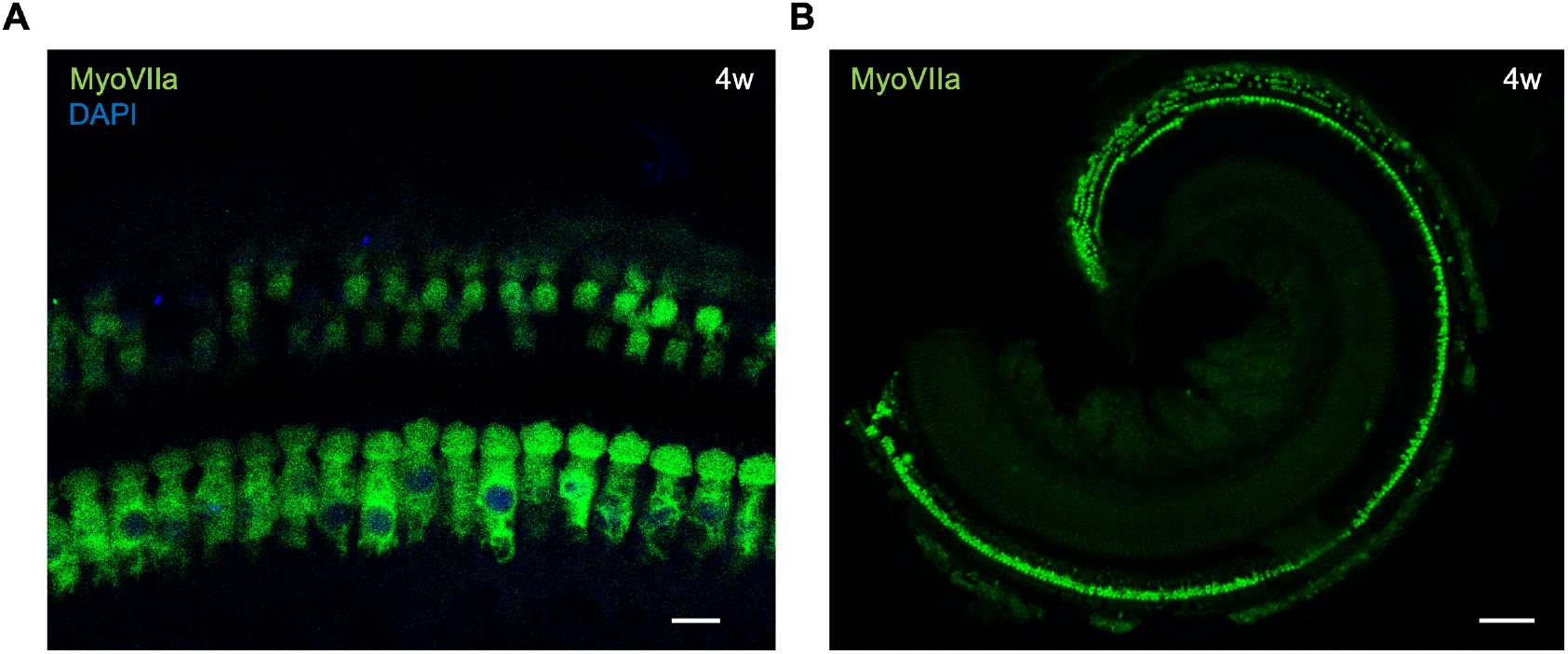
A. Whole-mount immunofluorescence of the 12kHz region from a 4w old *Syne4*^*-/-*^ mouse injected with AAV.Syne4^WT^. Hair cells labeled by MyoVIIa (green) and nuclei by DAPI (blue). C. Tile scan of a whole-mount immunofluorescence of a 4w old *Syne4*^*+/+*^ mouse injected with AAV.Syne4^AA^. Hair cells labeled by MyoVIIa (green). Scale bars = 10 and 100 μm for A and B, respectively.

**Supplementary Figure 3.**
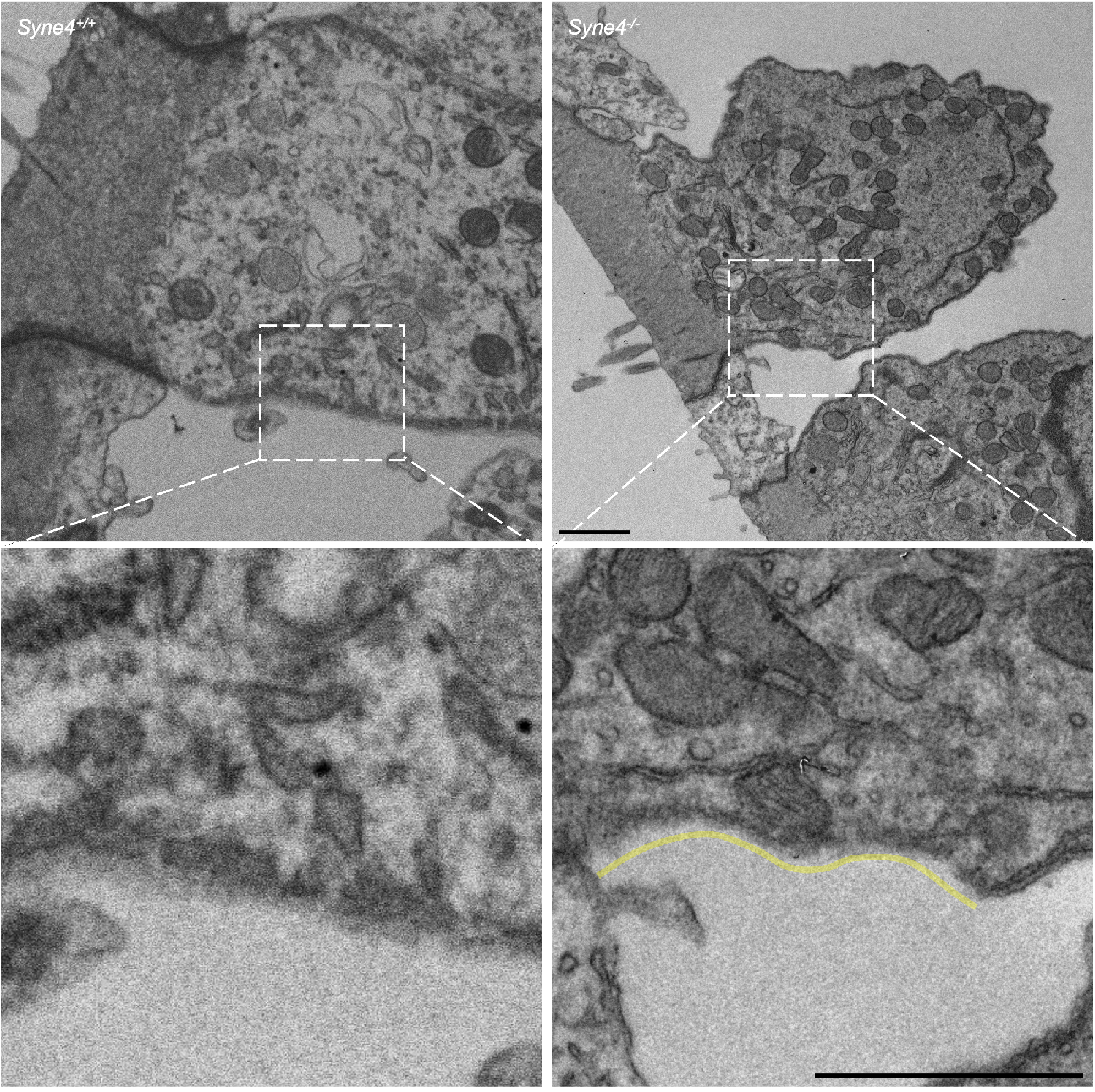
TEM of *Syne4*^*-/-*^ and *Syne4*^*+/+*^ OHC with insets showing high magnification of submembrane cisternae defects in *Syne4*^*-/-*^ OHC. Yellow line denotes cisternae defect. Scale bars = 1 μm.

**Supplementary Figure 4.**
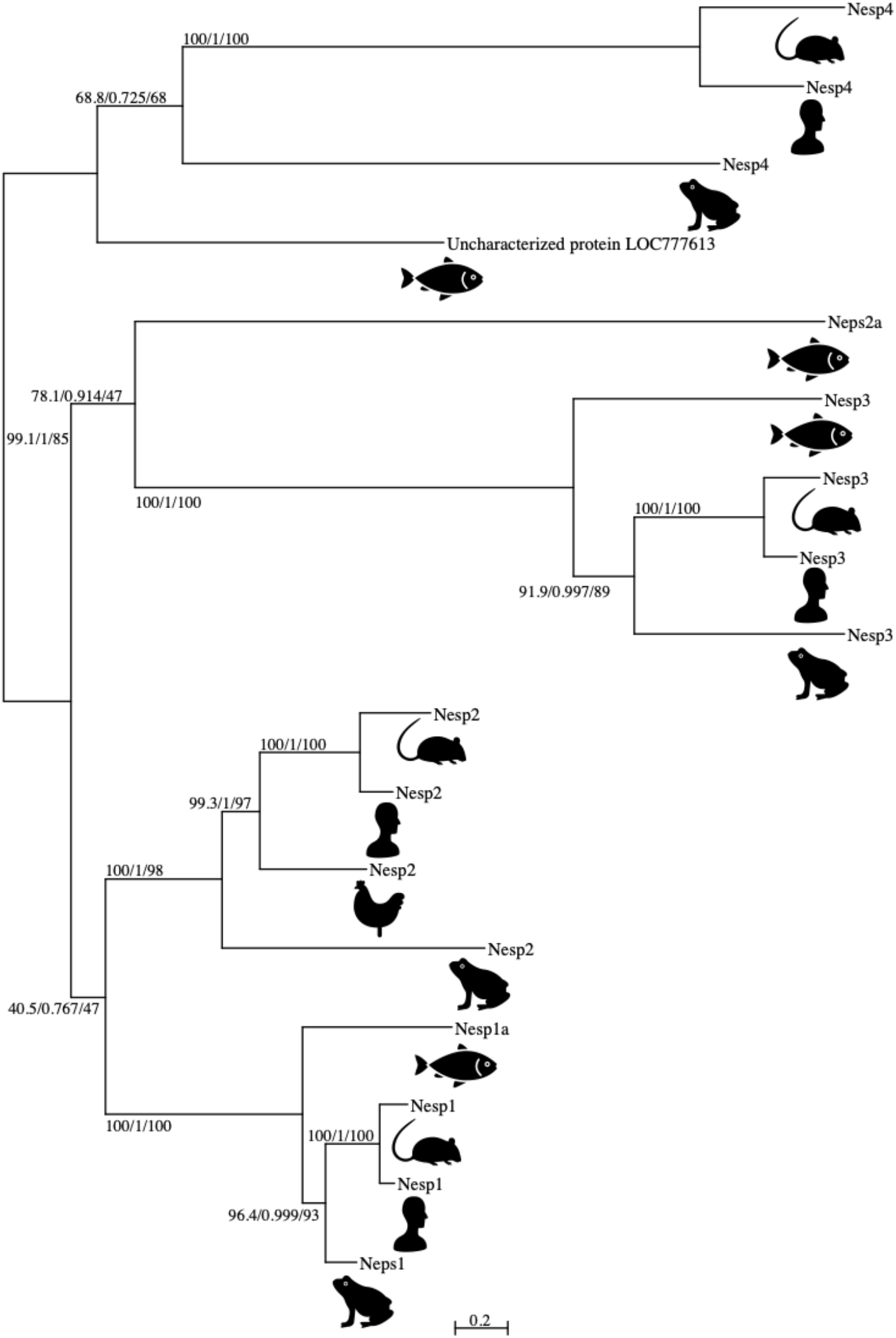
Phylogenetic analysis of nesprin proteins shows that nesprin-4 is conserved in vertebrates but is absent in invertebrates. Humanoid = *Homo sapiens*, rodent = *Mus musculus*, frog = *Xenopus tropicalis*, fish = *Danio rerio*, chicken = *Gallus gallus*. Units denote substitutions per site. Numbers next to each branch represent SH-Like, approximate Bayesian and ultrafast bootstrap support values, respectively.

